# Transient brain activity dynamics discriminate levels of consciousness during anesthesia

**DOI:** 10.1101/2023.10.09.560209

**Authors:** Scott Ensel, Lynn Uhrig, Ayberk Ozkirli, Guylaine Hoffner, Jordy Tasserie, Stanislas Dehaene, Dimitri Van De Ville, Béchir Jarraya, Elvira Pirondini

**Author notes:** Corresponding author*: Dr. Elvira Pirondini. co-senior authors (Béchir Jarraya, Elvira Pirondini).

## Abstract

The awake mammalian brain is functionally organized in terms of large-scale distributed networks that are constantly interacting. Loss of consciousness might disrupt this temporal organization leaving patients unresponsive. We hypothesized that characterizing brain activity in terms of transient events may provide a signature of consciousness. For this, we analyzed temporal dynamics of spatiotemporally overlapping functional networks obtained from fMRI transient activity across different states of consciousness. We first show a striking homology in spatial organization of networks between monkeys and humans, indicating cross-species similarities in resting-state fMRI structure. We then tracked how network organization shifts under different anesthesia conditions in macaque monkeys. While the spatial aspect of the networks was preserved, their temporal dynamics were highly affected by anesthesia. Networks expressed for longer durations and co-activated in an anesthetic-specific configuration. Additionally, hierarchical brain organization was disrupted with a consciousness-level- signature role of the default mode network. In conclusion, network temporal dynamics is a reliable and robust cortical signature of consciousness, paving the way to its clinical translation.

## Introduction

Recordings of spontaneous brain activity, assessed by resting-state functional magnetic resonance imaging (rs-fMRI), have provided key insights into the rich temporal dynamic and spatial organization of the awake brain [1–6]. Large-scale distributed networks can be extracted from spontaneous fluctuations of the fMRI signals over time. The spatiotemporal organization of these networks can reflect ongoing cognitive efforts and reflect changes in pathological conditions [7]. Recent studies suggested that the temporal dynamics of these networks could inform about the level of consciousness [8, 9].

Detecting residual consciousness in patients remains an open clinical problem. Several brain-imaging tests have been proposed to uncover residual signs of consciousness, but the majority of them require subjects to perform difficult and active cognitive tasks [10, 11]. Rs-fMRI could overcome these challenges and large-scale resting-state networks could provide reliable markers of the presence or absence of consciousness. With these premises, rs-fMRI dynamic connectivity has been investigated in sleep [3, 12, 13], anesthesia in humans [14, 15] and animals [1, 4, 16], and more recently in unresponsive or minimally conscious patients [8]. Interestingly, all these unconscious states shared hallmark brain changes. Specifically, cortical long-range interactions are disrupted in both space and time and spontaneous network-to-network transitions are less probable as configurations are rigid and tied to the underlying anatomical connectivity. By contrast, wakefulness has been associated with greater global integration and interareal cross-talk and a more flexible repertoire of functional brain configurations departing from anatomical constraints [1, 2].

While the dominance of rigid configurations tied to the underlying structural backbone constitutes a putative common signature of unconsciousness, lack of responsiveness can be associated with a variety of brain lesions, varying levels of vigilance, and distinct cognitive levels. We postulate that these differences could be reflected in the spatiotemporal reorganization of specific resting-state networks and brain regions. In this regard, previous studies reported a reduced activation of the visual and of the frontoparietal networks as well as the posterior cingulate cortex [17] and of the thalamus. Yet these characterizations remained limited and speculative hampering our capacity to distinguish level of consciousness.

A critical factor hindering the identification of brain network signatures of consciousness is the use of functional connectivity (FC) measures that use spatial and temporal information as mutually dependent. Indeed, the entirety of previous works used a sliding-window technique, where time courses from sets of brain regions are segmented into successive temporal windows and the different assessments of FC are applied to obtain time-evolving connectivity matrices [1, 2]. As a result, the dynamic evolution of functional networks interacting over time could not be fully captured and their properties remained singular in time. Here we overcome these limitations by deploying a recently proposed method termed innovation-driven co-activation patterns (iCAPs). iCAPs captures transients (i.e., moments of significant physiological changes in regional activation) [18–20] through regularized hemodynamic deconvolution of fMRI time series (i.e., framework of total activation, TA) and subsequent clustering of recurrent spatial patterns of transient activity into large-scale brain networks that can be both spatially and temporally overlapping.

Specifically, we adopted the iCAPs framework to recover functional brain networks from fMRI data recorded during different levels of anesthesia-induced unconsciousness in monkeys. For this, we first modified the deconvolution step to account for the hemodynamic response function due to monocrystalline iron oxide nanoparticles (MION). We then validated the translational properties of the identified large-scale brain networks comparing their spatial organization with networks obtained from human rs-fMRI. Finally, we computed temporal properties of these networks and showed that lack of consciousness resulted in longer duration and increased co-occurrence of networks paralleling previously reported reduced brain dynamics. Importantly, the networks co-occurrence and hierarchical organization differed from the conscious wakefulness in an anesthetic-specific configuration with an anesthetic-signature role of the default mode network.

## Results

We scanned 5 rhesus macaques during awake and under anesthesia with three different anesthetics (ketamine, propofol, and sevoflurane) and two different levels of consciousness (see **Supplementary Table 1** for a summary of the acquisitions). We used monkey sedation scale and electroencephalography traces to identify the level of consciousness for the different anesthetics. In all sessions and animals, the moderate anesthesia level for moderate propofol and moderate sevoflurane corresponded to level 3 over 4 on the monkey sedation scale; whereas the anesthesia level for ketamine, deep propofol, and deep sevoflurane corresponded to level 4 [2]. We then applied TA to the fMRI time series, that is hemodynamic deconvolution based on regularization of the temporal derivative of the deconvolved signal (**Supplementary Figure 1**), to obtain transients (i.e. moments of activity changes). Since a MION contrast agent was used during functional scans to increase functional sensitivity [1, 2], we integrated the MION hemodynamic response function (HRF) into the TA pipeline (**Supplementary Figure 2**). Significant innovation frames (i.e., spatial patterns of transient activity) were then obtained using a two-step thresholding process and subsequent K-means clustering procedure over all animals to label timepoints and obtain centroids that correspond to large-scale brain networks (i.e., iCAPs) that are potentially spatially overlapping. The optimal number of iCAPs was selected via consensus clustering. Finally, the backfitting of these innovation-driven patterns to the activity-inducing activity revealed time series that could be temporally overlapping (**Supplementary Figure 1**).

### Typical human large-scale networks were preserved in monkeys

In order to validate the translational potential of our results to human subjects, we first visually compared spatial networks extracted using the iCAP framework between humans and monkeys. We considered 7 publications [3, 20–25] that computed human iCAPs to gather the predominant human large-scale networks (**Supplementary Table 2**). We found that generally humans had an optimal number of clusters K = 18 ± 1.5, whereas monkeys had a reduced number of networks (K = 11, **Figure 1a**). Interestingly, humans and monkeys shared low-level functional networks such as the anterior and posterior cerebellum and primary and secondary visual networks (**Figure 1b**). Importantly, also anterior and posterior default mode networks were preserved in monkeys. In contrast, monkeys missed higher level cognitive networks such as attention, language, anterior salience and visuospatial networks. The lack of these high-level cognitive networks might explain the difference in optimal number of clusters. Yet, the majority of networks were preserved supporting the translational value of our results and justifying the animal-model choice in our work.

**Fig. 1.**
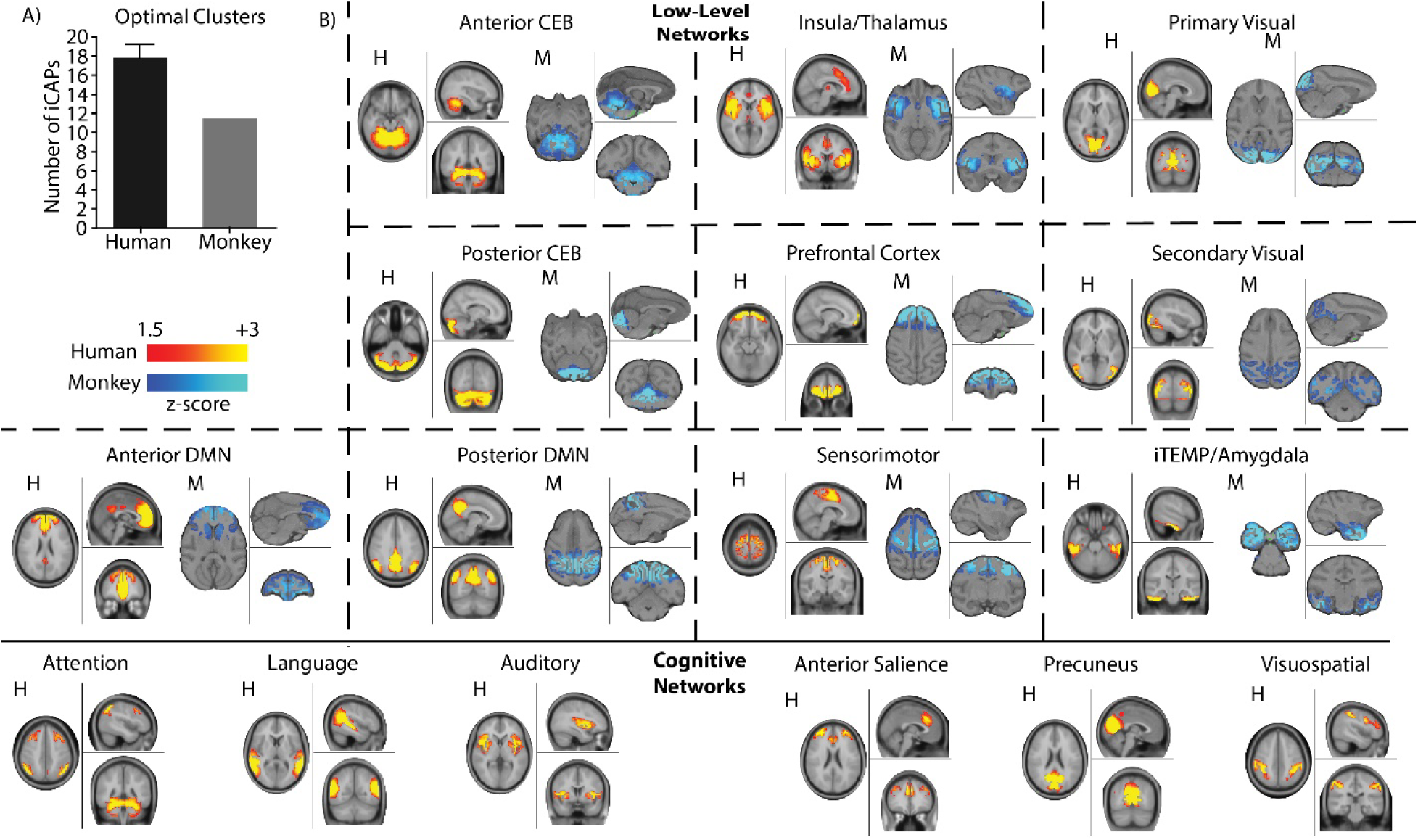
Humans and Monkeys have similar functional networks. **A)** Mean number of optimal clusters (iCAPs) found using consensus clustering for 7 studies in humans [3, 20–25] (black) and our monkey study (gray). Error bars represent SD over the 7 studies. **B)** *Top rows:* spatial patterns for the most common iCAP spatial clusters aggregated from previous studies in humans (red) compared to iCAP spatial clusters found from the monkeys (blue). All panels show a representative human network, marked with H, on the left, compared to a representative monkey network, marked with M, on the right. *Bottom row:* high-level cognitive networks that were found only in human studies. For all panels, colorbars represent the z-scored voxel values in each spatial iCAP.

### Slower brain dynamics during unconsciousness

We then compared the number of significant innovation frames between awake and anesthetic conditions. We observed a statistically significant difference between the number of significant positive and negative innovation frames in the awake condition as compared to all anesthetics (**Figure 2a**, mean ± SEM over animals for awake: 213 ± 16.9, 222 ± 18.8, ketamine: 145 ± 5.0, 138 ± 3.4, moderate propofol: 132 ± 4.3, 139 ± 5.8, deep propofol: 130 ± 6.0, 132 ± 3.0, moderate sevoflurane: 102 ± 3.2, 92 ± 4.9, deep sevoflurane: 96 ± 10.3, 84 ± 12.2 for significant positive and negative innovation frames, respectively). These significant innovation frames represent large changes in fMRI activations, and the statistically significant difference in their number highlights reduced brain dynamics when the animals were anesthetized.

**Fig. 2.**
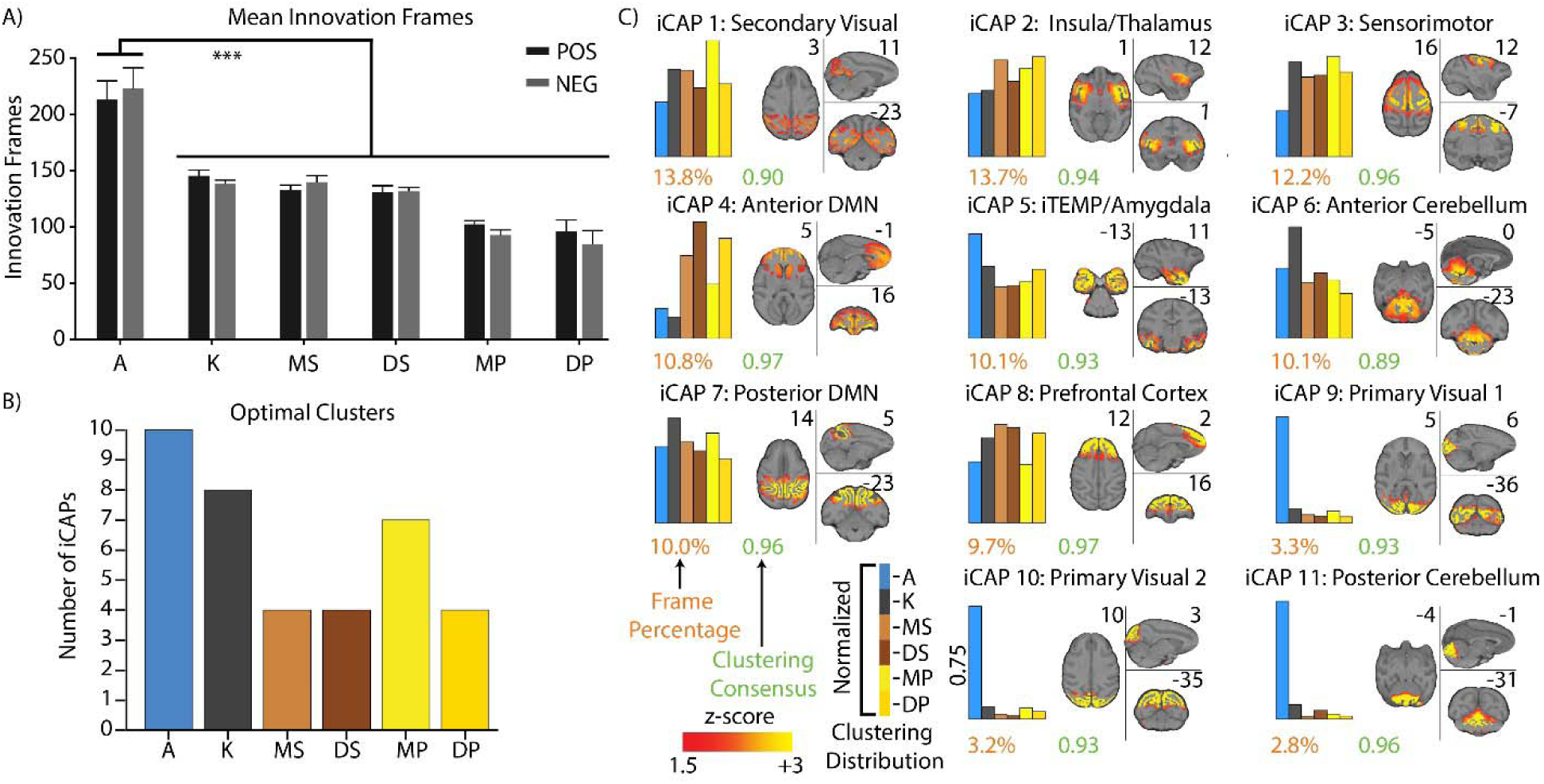
Clustering Distributions of Significant Innovation Frames. **A)** Bar graph representing the mean number of positive (black) and negative (grey) significant innovation frames for each condition. The error bars represent SEM over each animal and inference on the mean differences is performed by bootstrapping, with n=10,000 bootstrap samples; *** indicates statistical significance of p<0.001 adjusted with Bonferroni correction. (A: awake; K: ketamine anesthesia; MS: moderate sevoflurane sedation; DS: deep sevoflurane anesthesia; MP: moderate propofol sedation; DP: deep propofol anesthesia). B) Bar plot representing the optimal number of clusters (iCAPs) found for each individual condition using consensus clustering. **C)** Spatial patterns for the iCAPs obtained clustering together significant innovation frames from all conditions. The iCAPs are numbered according to the percentage of significant innovation frames that contributed to the recovery of that network (descending order). The number of significant innovation frames for each iCAP is shown below the spatial maps in orange font. The cluster consensus of each iCAP is reported in green. For each iCAP, the bar plots indicate the distribution of the significant innovation frames for each condition normalized over the different conditions. MNI coordinates of each brain slice are indicated in black close to each brain. The names of each iCAP are derived according to their correspondence with the CHARM and SARM atlas [26–28], which are presented in **Table 1**. In **Supplementary Figure 4**, the spatial patterns for the iCAPs extracted clustering the significant innovation frames for each condition separately.

These results were further validated by the optimal number of clusters. Indeed, when extracting iCAPs for each anesthetized and awake condition separately, we found that awake had the highest number of clusters (i.e. K=10), whereas the anesthetized conditions ranged from 8 to 4 clusters (**Figure 2b**).

### Spatial patterns of functional networks were preserved during unconsciousness

When clustering significant innovation frames from all conditions together, we obtained 11 large-scale iCAPs (**Supplementary Figure 3** for the results of the consensus clustering), representing the different functional networks that dominate brain activity across wakefulness and unconsciousness (**Figure 2c** and **Table 1**). The iCAPs corresponded to well-known functional large-scale networks obtained both in humans and monkeys with similar and different analysis approaches.

Specifically, the iCAPs included sensory-related networks such as the secondary visual system related to V2 and V3 brain areas, and part of the thalamus (iCAP 1), the primary visual system (iCAPs 9 and 10), and the sensory-motor system with a strong activation of the primary motor cortex (iCAP 3). Importantly, iCAP 1 was found to be equally prevalent across each condition; whereas, iCAPs 9 and 10 were predominantly active during the awake condition. The absence of a network for primary visual areas during unconsciousness parallels previous results on sleep in humans [3].

The anterior default mode network (aDMN, iCAP 4) and posterior DMN (pDMN, iCAP 7), instead, showed a similar activation during anesthetized conditions compared to the awake condition. It is of note that the anterior and posterior DMN were separately extracted, which emphasizes a dissociation between the temporal dynamics of these two regions (**Figure 3a**). This dissociation of the DMN into subnetworks is commonly observed in other iCAP studies supporting the reliability and robustness of the iCAP framework to extract large-scale brain networks [3]. A similar dissociation was also found for the cerebellum, which split into anterior (iCAP 6) and posterior (iCAP 11) cerebellum (**Figure 3b**). Importantly, the anterior portion was found across all conditions, while the posterior cerebellum was present almost exclusively in the awake condition again paralleling previous results on sleep in humans [1].

**Fig. 3.**
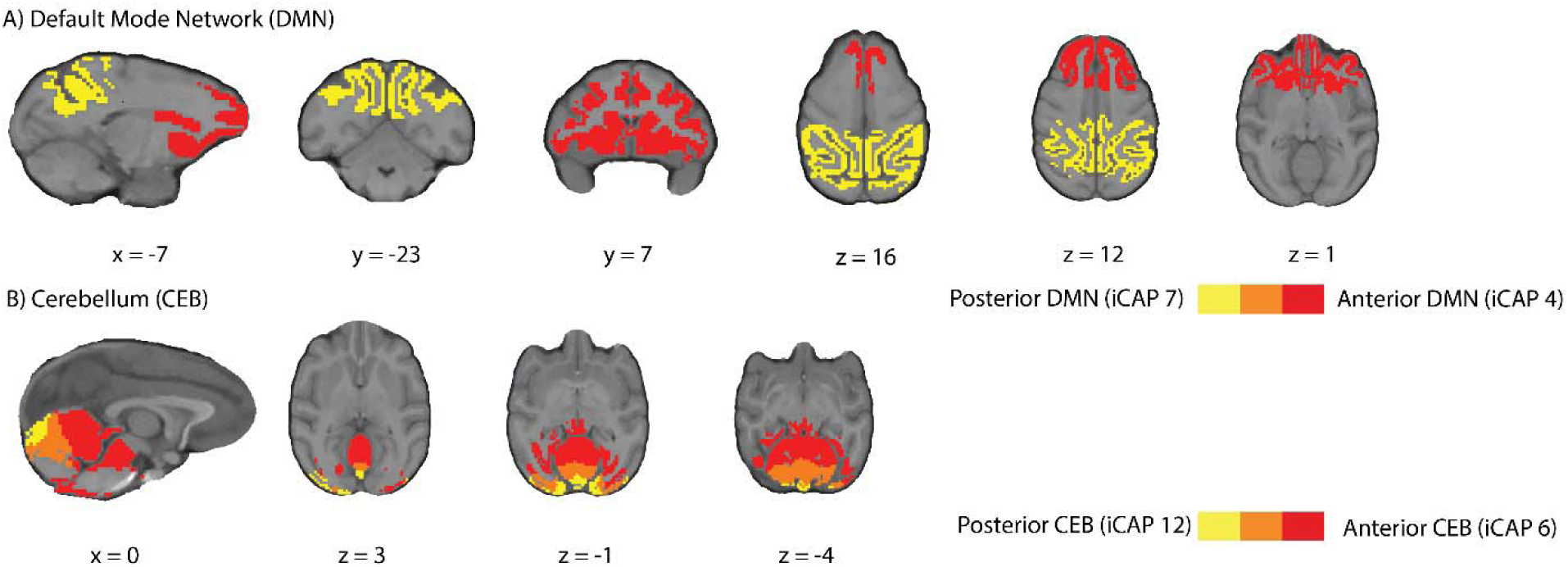
Dissociation of DMN and Cerebellum into posterior and anterior parts. **A)** Overlay of the iCAPs corresponding to the anterior DMN (red, iCAP 4) and posterior DMN (yellow, iCAP 7) with overlap (orange). **B)** Overlay of the iCAPs corresponding to the anterior cerebellum (red, iCAP 6) and posterior cerebellum (yellow, iCAP 7) with overlap (orange).

Lastly, iCAP 5, iCAP 8, and iCAP 2 represented the inferior temporal/amygdala (iTEMP/amygdala), prefrontal cortex (PFC), and insula network with portions of the thalamus and brainstem, respectively. The iTEMP/amygdala mostly resulted from significant innovation frames obtained during the awake condition, while iCAP 8 and iCAP 2 were composed of significant innovation frames equally distributed across all conditions.

Importantly, spatial patterns of the networks obtained when applying the iCAP framework to each condition separately matched those of the clusters found when conditions were combined (**Supplementary Figure 4**) further supporting that the spatial organization of the networks was preserved during unconsciousness.

### Altered temporal patterns of functional networks revealed consciousness-dependent brain dynamics

We then computed temporal metrics (in contrast to significant innovation frames) from the time series of each iCAP (**Supplementary Figure 1**). Analysis was limited to the first eight iCAPs as the remaining three (iCAPs 9-11) were not present during anesthesia.

Interestingly, iCAPs total duration was significantly longer in each of the anesthetized conditions compared to awake (**Figure 4a**) in particular for the secondary visual cortex (iCAP 1), insula (iCAP 2), anterior DMN (iCAP 4), and prefrontal cortex. Yet, iCAPs duration was comparable across anesthetics and levels of consciousness. We then looked at average duration of iCAPs in a similar fashion (**Figure 4b**). Indeed, while total duration only considers the sporadic activation of a network, average duration focuses on the length of continuous activation of each iCAP. Importantly, the continuous activation of iCAPs during anesthesia was also significantly greater than that in the awake condition for all iCAPs. This was consistent with the decrease in number of significant innovation frames observed in anesthetized conditions, highlighting the direct relation between slower dynamics and increased duration.

**Fig. 4.**
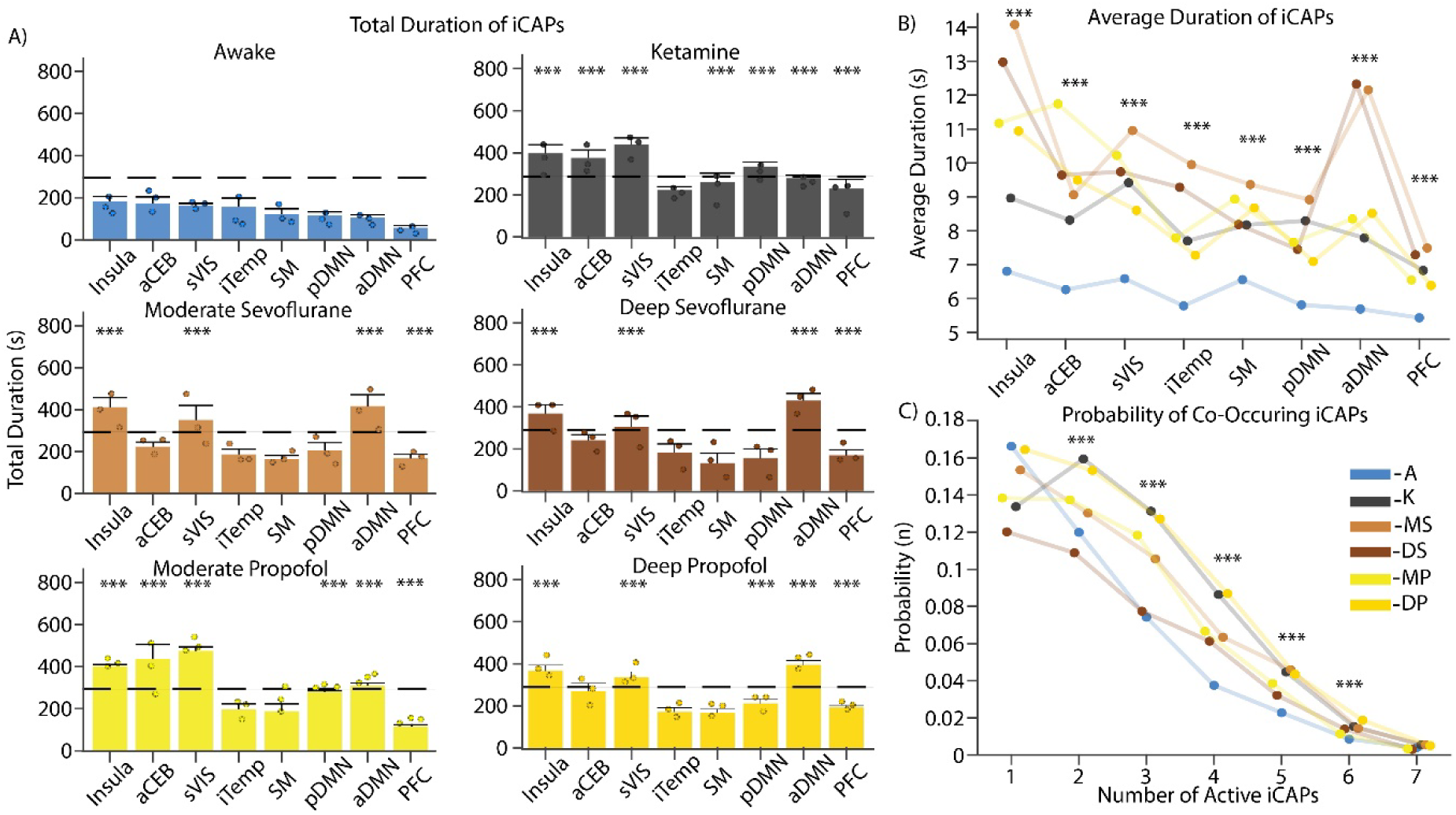
Durations of iCAPs in different conditions and probability of co-occurrences. **A)** Total duration of the first 8 iCAPs (in seconds) for each condition in descending order represented in awake. Each condition is represented in its own plot with awake (blue), ketamine (gray), moderate sevoflurane (light brown), deep sevoflurane (dark brown), moderate propofol (light yellow) and deep propofol (dark yellow). Error bars on each bar represent SEM over each animal and the black dashed line represents 25% of total possible duration. Dots represent average values for each animal individually. **B)** Average durations (in seconds) of iCAPs reflecting the length of continuous activity**. C)** The probability of a different number of iCAPs to overlap is displayed for each condition. For all panels, inference on the mean differences is performed by bootstrapping, with n=10,000 bootstrap samples; *** indicates statistical significance of p<0.001 adjusted with Bonferroni correction.

To explore beyond the iCAPs’ individual temporal properties, we then evaluated the temporal overlap between iCAPs and computed the probability of having one, two or more iCAPs occurring concurrently (**Figure 4c**). iCAPs in the awake condition had a higher probability of occurring alone, whereas during anesthesia they had a higher probability of co-occurring. These results are consistent with iCAPs’ individual temporal properties. Indeed, the longer total and average durations of iCAPs in anesthetized conditions resulted in an increased co-occurrence of iCAPs over time.

In summary, iCAPs temporal properties further supported reduced brain dynamics during unconsciousness that was evident in longer duration and increased co-occurrence of brain networks but with no differentiation across anesthetics.

### Networks co-occurrence revealed anesthetic-specific brain dynamics

To further explore the dynamic interactions across networks, we computed the temporal co-occurrence for every pair of iCAPs (**Figure 5a**). As expected, anesthetized conditions had stronger co-occurrence between pairs of iCAPs when compared to the awake condition. Some pairs were shared across anesthetics, whereas others were anesthetic-specific. Indeed, every anesthetic, except ketamine, had a significantly increased co-occurrence for secondary visual cortex (iCAP 1) and insula (iCAP 2) with anterior DMN (iCAP 4) and for secondary visual cortex and insula. Similarly, secondary visual cortex and insula co-occurred with prefrontal cortex (iCAP 8) for ketamine and sevoflurane. Additionally, while the anterior DMN co-occurred with the anterior cerebellum (iCAP 6) for sevoflurane and propofol, the posterior DMN (iCAP 7) co-occurred with the anterior cerebellum for ketamine and propofol. The anterior DMN co-occurred also with the posterior DMN for propofol and with the iTEMP/amygdala (iCAP 5) for sevoflurane. Finally, ketamine had a significantly increased co-occurrence for prefrontal cortex with the posterior DMN and the anterior cerebellum.

**Fig. 5.**
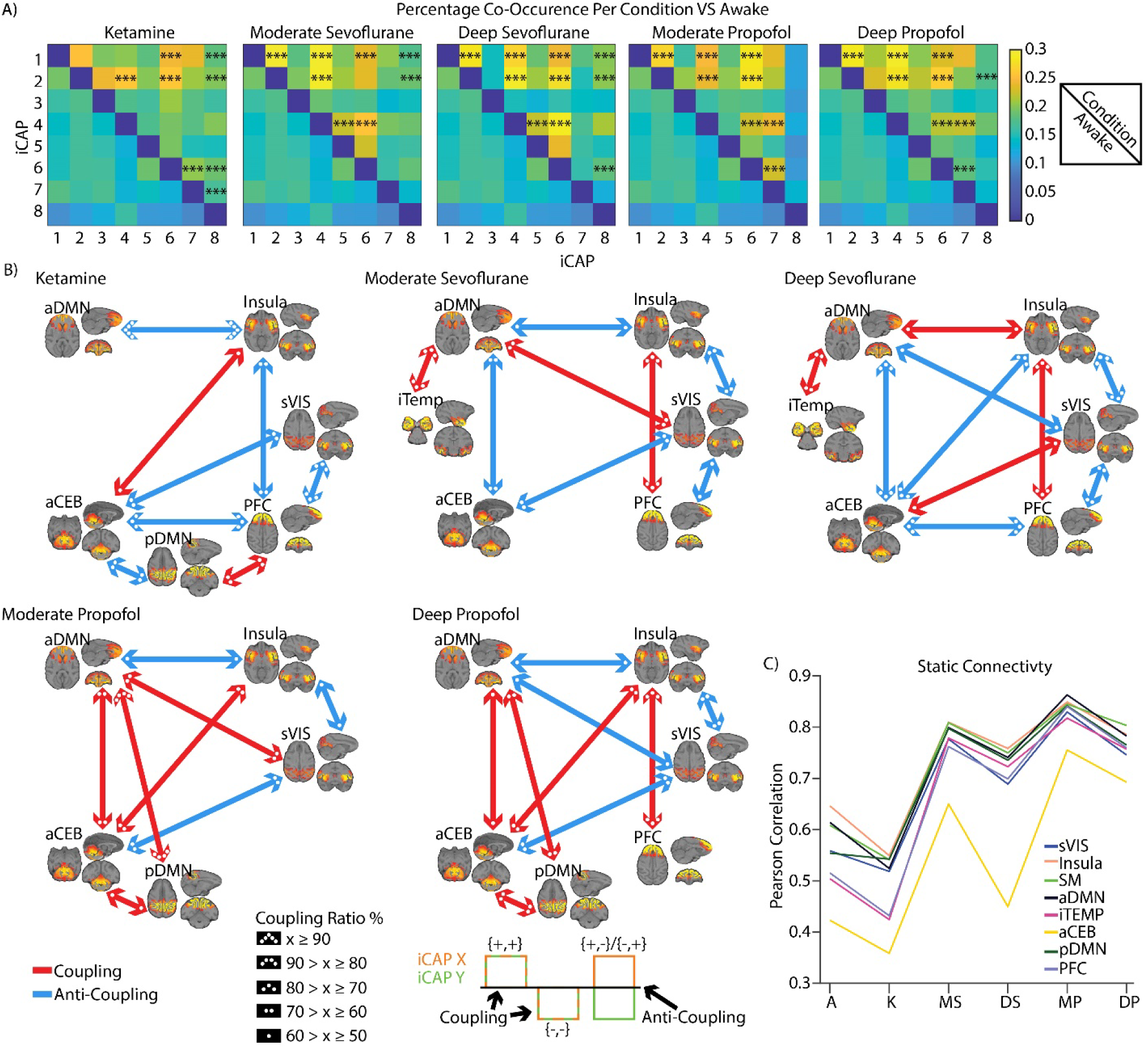
Pairwise coupling between iCAPs across different conditions compared to awake. **A)** The co-occurrence pertains to the number of times in a pair of iCAPs both iCAPs were active, divided by the total number of time points where at least one of them was active. This is computed together for same or opposite signed activations. Top right of each square is a sedative condition and the bottom left of each square is the awake condition. Inference on the mean differences is performed by bootstrapping, with n=10,000 bootstrap samples; *** indicates statistical significance of p<0.001 adjusted with Bonferroni correction. **B)** Significant total co-occurrences are broken down into coupling and anti-coupling. Coupling and anti-coupling are indicated in red and blue, respectively, and the degree of coupling or anti-coupling is indicated by the number of dots in each arrow head. **C)** Mean of pairwise Pearson correlation applied to the iCAP time courses. Each color indicates a different iCAP.

Because iCAPs could have a positive or negative activation, we then considered co-occurring activity in terms of polarity and we termed coupling when two iCAPs were both positively or negatively activated [(+,+),(-,-)] and anti-coupling when the two iCAPs had opposite activation [(+,-),(-,+)] (**Figure 5b**). Not surprisingly, some couplings and anti-couplings were common across anesthetics. Specifically, the insula coupled with the prefrontal cortex for all the conditions except moderate propofol and anti-coupled with the anterior DMN, except in deep sevoflurane (coupling). The secondary visual cortex, instead, anti-coupled with the insula except for deep sevoflurane (i.e., deepest level of unconsciousness). Interestingly, the coupling between the secondary visual cortex and the anterior DMN was positive for light levels of consciousness (moderate propofol and moderate sevoflurane) and negative for deeper levels of consciousness (deep propofol and sevoflurane). Under ketamine, moderate, and deep sevoflurane the prefrontal cortex was anti-coupled with the secondary visual cortex and the anterior cerebellum (not for moderate sevoflurane) showing similar interactions across these anesthetics in particular during deep anesthesia. However, ketamine was distinct from the other anesthetics as it had a higher proportion of anti-coupled pairs representing a strong difference in brain dynamics. Finally, the anterior DMN and anterior cerebellum were positively coupled under propofol and negatively under sevoflurane.

Importantly, couplings and anti-couplings between iCAP pairs paralleled findings obtained with a classic functional connectivity (FC) analysis. Indeed, we computed Pearson correlation of the iCAPs time courses and we found that pairwise FC across iCAPs increased in all the anesthetics as compared to awake except for ketamine (**Figure 5c**). Ketamine’s reduced correlation between networks matched its stronger anti-coupling of iCAPs. Interestingly, the anterior cerebellum was unique as compared to other iCAPs showing a reduced connectivity, which parallels its susceptibility to be anti-coupled with other networks.

In summary, while generally iCAPs co-occurred more during unconsciousness for all anesthetics, different drugs and more importantly different levels of anesthetic induced unconsciousness increased coupling and anti-coupling between specific iCAPs, supporting the need to assess spatiotemporal networks interaction to discern alertness levels.

### Network hierarchical organization revealed an anesthetic-specific role of the DMN

In order to reveal a hierarchical organization across large-scale networks, we expanded our analysis to all possible combinations of iCAPs. We then applied hierarchical clustering of iCAPs using the observed combinations as features (**Figure 6**). The dendrogram reflects a hierarchy of iCAPs based on their frequency of occurrence together. In awake, similarly to previous results in humans [20], iCAPs were grouped into two large clusters of sensory and default-mode networks. Interestingly, this division was not preserved during anesthesia. Indeed, except for ketamine, the anterior and posterior DMN split into two different clusters further supporting the strong dynamical dissociation of the DMN into subnetworks, which can be captured only with the iCAP framework. In the anesthesia conditions, except for ketamine, the anterior DMN was clustered with the secondary visual, insula, and anterior cerebellum. Instead, in the other large branch, there was a consistent organization of the posterior DMN, prefrontal cortex, iTEMP/amygdala, and sensory-motor cortex. Within this split light and deep propofol and moderate sevoflurane shared the same hierarchical organization as opposite to deep sevoflurane. Ketamine, instead, was unique as it shared the same organization of this split but with included the anterior DMN.

**Fig. 6.**
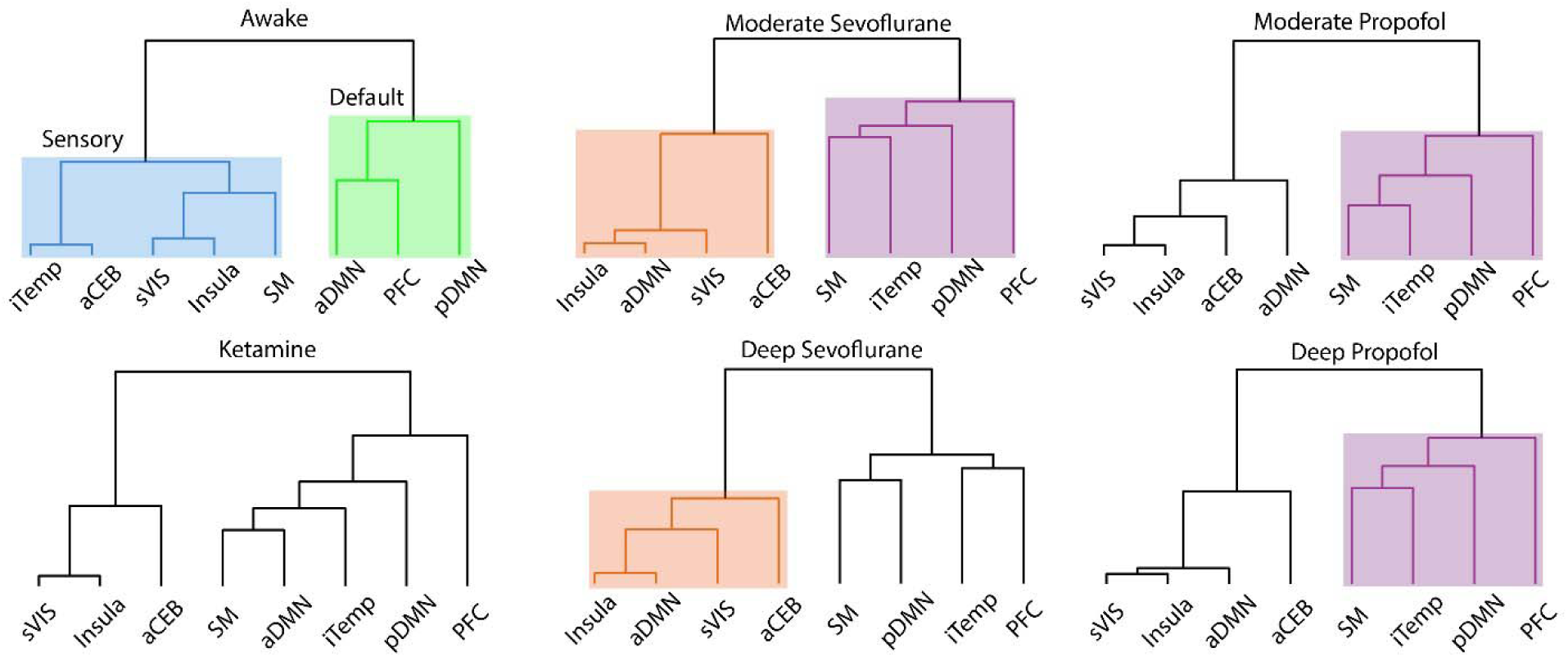
Hierarchical clustering of iCAPs according to their temporal overlap. The dendrogram minimizes the distance at each leaf with respect to the neighboring leaf, clustering the most similar iCAPs. Each condition is represented by their own tree with some branches highlighted by unique colors to emphasize similarities or differences for each cluster. Awake is the only condition separated into a sensory and default networks where the other conditions have unique formations.

In summary, unconsciousness disrupts the hierarchical organization of brain dynamics dissociating the anterior and posterior DMN. Importantly, the association of the latter with other networks appears to be pivotal in differentiating anesthetics and levels of consciousness.

## Discussion

Recent experiments in sedated humans, rats, and monkeys [14, 15, 29] have shown that under general anesthesia spontaneous brain activity converges to a few or even a single dominant brain dynamical pattern. This reduced dynamics was confirmed in unresponsive or minimally conscious patients [8] validating the use of pharmacologically induced anesthesia models to study loss of consciousness. However, these decreased dynamics may not simply derive from an overall reduction of functional connectivity, but it could be the result of a complex orchestration of preserved and suppressed functional networks [30]. This composition may be occulted when using static FC or sliding window methods [1, 2], challenging our ability to distinguish levels of consciousness. Here we overcome the limitations imposed by static FC or sliding windows methods by clustering moments of significantly changing brain activity from multiple fMRI sessions while animals at rest were awake or under anesthesia to extract large scale brain networks or iCAPs. The use of transient fMRI activity allowed us to obtain temporally overlapping spatial networks and to compute their time courses at a time-scale of seconds. We established that functional brain networks preserved spatial organization in unconscious states; yet they had fewer dynamic fluctuations than conscious states, which resulted in longer temporal activation and more probable network co-activations. Importantly, network co-occurrence showed anesthetic-specific trends and a hierarchical organization specific to the level of consciousness. Altogether, these results advance the use of rs-fMRI to detect the presence or absence of consciousness.

Here we discuss our findings with an emphasis on key anatomical brain structures and temporal measures that distinguish different anesthetics and level of consciousness.

### A whole-brain dynamic alteration

During unconsciousness, the amount of transient brain activity significantly decreased as compared to awake resulting in a reduced number of clusters, which was explained by the absence of activation of primary visual cortex and posterior cerebellum. The lack of a primary visual cortex network might be explained by the difference in eyes-opened during awake, and eyes-closed when under anesthesia induced by a muscle-blocking agent. The absence of the posterior, but not the anterior, cerebellum, instead, might be explained by the organization of the cerebellum lobules. Indeed, the anterior lobules have been reported to be responsible for the sensorimotor domain, while the posterior ones for the cognitive domain [31]. Additionally, previous works reported a lack of the posterior cerebellum network during sleep [3] and a marked correlation of larger lobules (i.e., anterior region of the cerebellum) [32] with slow-wave sleep and slow spindles in different sleep stages [33]. Importantly, the spatial organization of the other brain networks was preserved.

To explain how the predominance of temporal reorganization can explain loss of consciousness during anesthesia while anatomical networks are preserved, we showed a global increase in simultaneous coupling and anti-coupling signaling instability of network synchronization and ineffective brain integration. These results are in line with previous studies [34] and commonly interpreted to reflect reduced consciousness during sleep [35]. We here showed that a similar mechanism might occur during anesthesia further supporting the idea that consciousness is not the persistence of functional brain networks, but rather the degree of interactions among them [34, 36–38]. This lack of integrity and synchronization could be due to a reduced activation of the thalamus. Yet, the role of the thalamus in inducing loss of consciousness remains controversial. Indeed, several studies show that anesthesia-induced loss of consciousness is mostly associated with a change in cortical correlation rather than with an alteration of thalamic activity [39, 40]. On the other hand, because of its pivotal role in the exchange of information between the periphery and cortex, the thalamus has been hypothesized to be essential in inducing loss of consciousness [41]. In this regards, recent studies showed that stimulation of the thalamus during anesthesia restored arousal and wake-like neural processing as depicted by brain fluctuations similar to the awake state [9, 42, 43]. The spatiotemporal organization of iCAPs further supports this second hypothesis. Indeed, both iCAPs 1 and 2, which included portions of the thalamus, presented the strongest temporal changes during loss of consciousness. Specifically, they fluctuated with the longest average durations and had an increased co-occurrence with other networks, particularly with the anterior DMN and the prefrontal cortex. These areas are highly coupled together through the lateral forebrain bundle which connects the temporo-parietal DMN nodes, whose connectivity is believed to be essential for conscious awareness [44, 45]. Interestingly, these cortical areas also had a significantly longer total duration during anesthesia further suggesting that loss of consciousness might induced by a reduced activation of the thalamus that caused a slowing down of the prefrontal cortex and DMN.

Importantly, while the reduced fluctuations in the thalamo-prefrontal pathways was a common signature of anesthesia-induced loss of consciousness no matter the initial molecular mechanism of the anesthetic, co-activations of other brain areas were, instead, anesthetic-specific. In particular the activation of the anterior cerebellum differentiated the most across opiates. While the cerebellum has long been considered a marginal target for general anesthesia [46], recent studies suggested a possible involvement of this structure explained by its strong connectivity with prefrontal and frontal cortical areas [47]. Our results show support for this hypothesis. Indeed, the anterior cerebellum anti-coupled with the anterior DMN when unconsciousness was induced with sevoflurane, but these networks positively coupled during propofol-induced anesthesia. The anterior cerebellum likewise anti-coupled more with the prefrontal cortex under ketamine and sevoflurane sedation than during the awake condition. Importantly, these more probable cortical co-activations of the anterior cerebellum during anesthesia as compared to the awake condition reflect a reduction of low-frequency fluctuations in the frontal regions and cerebellum, which is consistent with frontal to sensory-motor cortical disconnection as a possible mechanism of loss of consciousness [48].

### The default mode network distinguishes level of consciousness

Also, the hierarchical organization of the networks was unique between conscious and unconscious states and differentiated levels of alertness. Indeed, brain activity during consciousness arranged into components related to sensory and attention, matching organization shown in previous studies of awake humans [3]. During unconsciousness, instead, this organization was altered in particular at the level of the DMN connectivity. Specifically, the anterior and posterior parts of the DMN split into two different hierarchical clusters: the anterior DMN reorganized with the secondary visual, insula, and anterior cerebellum; while the posterior DMN clustered with the prefrontal cortex, iTEMP/amygdala and sensory-motor cortex. These findings parallel previous works showing that functional correlation decreased in the DMN during all the anesthetic conditions, as compared to the awake condition [49–51]. Similarly, the DMN dissociates into posterior and anterior parts upon reaching deep sleep [38, 52] with an increased activation of the posterior DMN [3]. These similarities suggest a common neurophysiological role of the posterior DMN in inducing unconsciousness between sleep and anesthesia. Importantly, within the hierarchical split containing the posterior DMN, moderate and deep propofol, and moderate sevoflurane shared the same hierarchical organization as opposite to deep sevoflurane. Finally, ketamine had a unique hierarchy of iCAPs with the anterior and posterior DMN in the same split together with the prefrontal cortex, iTEMP/amygdala and sensory-motor cortex. If we assume a finer definition of level of alertness with moderate propofol the lowest level of unconsciousness and deep sevoflurane and ketamine the strongest, our results hint to a critical role of the posterior DMN; i.e., the posterior cingulate cortex, in determining the level of consciousness.

### Clinical translation

Importantly, previous works in humans using the iCAP method reported large-scale brain networks that are consistent with our results demonstrating similar cognitive processes and neurobiological correlates in humans and non-human primates which enables translation of discoveries. Common FC network structures across species were reported also in other studies further supporting the generalizability of our work to patients [4, 29, 53–57]. Importantly, the similarity across networks was preserved even if our approach uses a custom MION HRF instead of the standard HRF. These similarities open up the possibility that residual consciousness could be monitored through the dynamics of brain activity and tentatively suggests new biomarkers of conscious activity. Our findings could help ameliorate the accurate diagnosis of patients with disorders of consciousness and further develop future generations of anesthesia monitoring devices.

## Conclusion

Dynamic functional connectivity for fMRI has been investigated in different sleep stages [3, 12, 13], under different anesthetic sedation, and more recently in unconscious patients [8] showing a decreased brain dynamic. However, the investigation regarding the affected brain areas remained limited and speculative, thus hindering our capacity to distinguish alertness. Here we deployed the iCAPs framework that has the unique ability to extract spatially and temporally overlapping functional networks. We observed a clear dissociation of both the DMN and cerebellum into anterior and posterior networks. Thanks to these features, we identified key brain areas responsible for loss of consciousness and with different effects depending on the anesthetic and the level of consciousness and involving the default mode network, the anterior cerebellum, and the thalamus. Specifically, the activation of the thalamus may play a central role, common to all anesthetics, on reducing the incoming information flow from the periphery that reduces the whole-brain dynamics. Simultaneously, a decreased activation of the anterior cerebellum causes a disconnection with the frontal regions, with an anesthetic-specific differentiation between the anterior DMN and the prefrontal cortex. Finally, the posterior DMN regulates the desynchronization of multiple networks inducing different level of consciousness.

## Materials and Methods

### Animals

5 rhesus macaques (Macaca mulatta), 1 male (monkey J) and 4 females (monkeys A, K, Ki, and R), 5 to 8 kg, 8 to 12 year of age, were tested; three for each arousal condition (awake: monkeys A, K, and J; propofol anesthesia: monkeys K, R, and J; ketamine anesthesia: monkeys K, R, and Ki; sevoflurane anesthesia: monkeys Ki, R, and J). See **Supplementary Table 1** for details on the exact number of acquisitions per animal and condition. All procedures were conducted in accordance with the European Convention for the Protection of Vertebrate Animals used for Experimental and Other Scientific Purposes (Directive 2010/63/EU) and the National Institutes of Health’s Guide for the Care and Use of Laboratory Animals. Animal studies were approved by the institutional Ethical Committee (Commissariat à l’Énergie atomique et aux Énergies alternatives; Fontenay aux Roses, France; protocols 10-003 and 12-086). Part of the data used in this work were already presented in [1, 2]. Therefore, anesthesia protocols were the same of these previous works.

### Anesthesia protocol

The monkeys were administered anesthesia using either ketamine [58], propofol [1, 58], or sevoflurane. We gauged the depth of anesthesia by employing the monkey sedation scale, which considers spontaneous movements and responses to various external stimuli such as shaking, prodding, toe pinching, and assessing the corneal reflex. Clinical scores were determined at the outset and conclusion of each scanning session, coupled with continuous electroencephalography monitoring.

For ketamine anesthesia, the monkeys received an initial intramuscular injection of ketamine (20 mg/kg; Virbac, France) for induction, followed by a continuous intravenous ketamine infusion (15 to 16 mg · kg^−1^ · h^−1^) to maintain anesthesia. Atropine (0.02 mg/kg intramuscularly; Aguettant, France) was administered 10 minutes prior to induction to reduce salivary and bronchial secretions. For propofol anesthesia, the monkeys were scanned at moderate propofol sedation and deep propofol anesthesia levels. For this, the animals were trained to receive an intravenous propofol bolus (5 to 7.5 mg/kg; Fresenius Kabi, France) for anesthesia induction, followed by target-controlled propofol infusion (moderate propofol sedation: 3.7 to 4.0 µg/ml; deep propofol anesthesia: 5.6 to 7.2 μg/ml) based on the “Paedfusor” pharmacokinetic model [59]. Despite the “Paedfusor” model being validated in humans, we previously applied it to macaque monkeys, observing stable clinical scores and electroencephalography activity during propofol anesthesia sessions [1, 58]. Finally, for sevoflurane anesthesia, the monkeys were subjected to moderate and deep sevoflurane anesthesia. The induction of anesthesia was done by intramuscular ketamine injection (20 mg/kg; Virbac), followed by sevoflurane administration (moderate sevoflurane anesthesia: sevoflurane inspiratory/expiratory, 2.2/2.1 volume percent; deep sevoflurane anesthesia: sevoflurane inspiratory/expiratory, 4.4/4.0 volume percent; Abbott, France). We waited a minimum of 80 minutes [60] to elapse before initiating scanning sessions during sevoflurane anesthesia to ensure the elimination of the initial ketamine injection. To prevent movement-related artifacts during magnetic resonance imaging acquisition, a muscle-blocking agent (Cisatracurium, 0.15 mg/kg bolus intravenously, followed by continuous intravenous infusion at a rate of 0.18 mg · kg^−1^ · h^−1^; GlaxoSmithKline, France) was co-administered during the ketamine and deep propofol sedation sessions.

In all anesthesia experiments, the monkeys were intubated and ventilated. Vital signs such as heart rate, noninvasive blood pressure (systolic/diastolic/mean), oxygen saturation, respiratory rate, end-tidal carbon dioxide, and cutaneous temperature were continuously monitored (Maglife, Schiller, France) and recorded online (Schiller).

### Electroencephalography

To assess the depth of anesthesia, we utilized scalp electroencephalography (EEG) with a magnetic resonance imaging (MRI) compatible system and custom-made caps, as detailed previously [1]. Our analysis was conducted in real-time through visual inspection of EEG patterns following established guidelines [58]. The awake condition (level 1) was characterized by posterior α waves (with eyes closed) and anterior β waves. For ketamine, sedation level was 4, which was characterized by intermittent polymorphic δ activity (0.5 to 2 Hz) of large amplitude, overlaid by low-amplitude β activity [61], and an increase in γ power (30 to 100 Hz) [62]. For propofol, sedation levels were 3 (moderate propofol sedation) featuring diffuse and wide α waves, along with anterior theta waves [63], and level 4 (deep propofol) marked by diffuse delta waves, low-amplitude waves [64], and anterior α waves (10 Hz) [65]. For sevoflurane, sedation levels were 3 (moderate sevoflurane) featured by elevated frontal delta, α, and β waves [66], and level 4 (deep sevoflurane anesthesia) characterized by diffuse delta waves and anterior α waves [66].

### Functional Magnetic Resonance Imaging Data Acquisition

The data were collected between July 2011 and August 2016. Monkeys were scanned on a 3-Tesla horizontal scanner (Siemens Tim Trio, Germany) with a single transmit-receiver surface customized coil. Each functional scan consisted of gradient-echoplanar whole-brain images (repetition time = 2,400 ms; echo time = 20 ms; 1.5-mm^3^ voxel size; 500 brain volumes per run). Before each scanning session, monocrystalline iron oxide nanoparticle (Feraheme, AMAG Pharmaceuticals, USA; 10 mg/kg, intravenous) was injected into the monkey’s saphenous vein [1]. For the awake condition, monkeys were implanted with a magnetic resonance compatible headpost and trained to sit in the sphinx position in a primate chair without performing any task [1, 67]. The eye position was monitored at 120 Hz (Iscan Inc., USA). For the anesthesia sessions, animals were positioned in a sphinx position, mechanically ventilated, and their physiologic parameters were monitored.

### Functional Magnetic Resonance Imaging Preprocessing

Functional images were preprocessed using the Pypreclin pipeline for monkey fMRI [68]. Images were slice-time corrected with FSL slice timer function (FMRIB’s Software Library – FSL, Oxford, U.K) [69]. B0 inhomogeneities and B1 field were corrected using the SyN function and N4 normalization of the Advanced Normalization Tool (ANTS). Images were reoriented, realigned, and rigidly co-registered to the anatomical template of the macaque Montreal Neurologic Institute (MNI, Montreal, Canada) space with use of JIP align (http://www.nmr.mgh.harvard.edu/~jbm/jip/, Joe Mandeville, Massachusetts General Hospital, Harvard University, MA, USA) and Oxford Centre Functional Magnetic Resonance Imaging of the Brain Software Library software (United Kingdom, http://www.fmrib.ox.ac.uk/fsl/; accessed February 4, 2018) [67]. The data were denoised using the non-human primate adapted ICD-FIX command (https://github.com/Washington-University/NHPPipelines) for spatial Independent Component Analysis (ICA, i.e. melodic) followed by automatic classification of components into ‘signal’ and ‘noise’. We then applied spatial smoothing using an isotropic Gaussian kernel of 3 mm full width at half maximum.

### Total Activation and the iCAPs framework

In order to extract large-scale brain networks and their temporal characteristics we deployed the iCAP pipeline [18, 20, 23, 24, 70, 71] (**Supplementary Figure S1**). We first implemented the Total Activation framework which applies MION informed deconvolution (**Supplementary Figure S1**) to the pre-processed fMRI time series of each run separately to reliably retrieve activity-inducing signals. More details about the implementation the TA framework can be found in Karahanoğlu et van De Ville, 2015 [20].

Then, for each activity-inducing time courses we removed the first ten volumes (i.e. 490 volume kept in total per run) and significant activation change-points (i.e. transients) were computed as the temporal derivative of these activity-inducing signals. After TA is ran, we extracted sparse activation points which were computed by taking the temporal derivative of the activity-inducing signals. Significant innovation frames (i.e. frames with significant transitioning activities) were selected with a two-step thresholding procedure with temporal and spatial thresholds selected based on previous work [18–21, 23, 24]. This two-step thresholding allowed to select only frames that contained significantly transient activity and to avoid including spurious connectivity patterns. The purpose of the temporal thresholding was, for each voxel, to find the time points where the activity was significantly high/low (i.e. positive/ negative). To determine the temporal threshold a surrogate data set, created by phase-randomization of the real data, was used to build a surrogate distribution where the lowest 1^st^ and highest 99^th^ percentile was used to select significant voxels. We did that for all voxels, and obtained, for each time point, a map of significant positive and negative transients (regions that are jointly activation, positive, and regions that jointly deactivation, negative). We then applied spatial thresholding on these maps, and only select those that have more than 5% of significant voxels (i.e., significant innovation frames).

Significant innovation frames across all animals, sessions, and anesthetics were then fed into a temporal k-means clustering to obtain large-scale resting state networks, the iCAPs. The optimum number of 11 clusters was determined by consensus clustering [72]. The method involved subsampling of the data and multiple runs of the clustering algorithm. The consistency of each frame grouped in a similar cluster was monitored through a consensus metric. The number of clusters was evaluated from K = 5 to 15.

We chose K = 11 to be the optimal number of clusters by looking at the trend of the area under the curve (AUC) corresponding to the consensus clustering matrices (**Supplementary Figure 3A**). We also looked at the slope of the cumulative distribution function (CDF) of the consensus clustering matrices (**Supplementary Figure 3B**). The optimal number also coincides with the K that has the highest cluster consensus (**Supplementary Figure 3C**).

Finally, session specific iCAP time courses were obtained by transient-informed spatiotemporal back-projection of the 11 spatial maps (i.e. the large-scale rs-fMRI networks extracted when clustering together significant innovation frames concatenated across all sessions) onto the individual activity-inducing signals. For each subject the iCAPs time courses were Z-scored and thresholded (|Z> 1.5) to highlight active and de-active time points. The choice of this particular threshold was motivated by previous works that implemented TA and iCAP framework [3, 21]. For each iCAP, and session we then computed the total duration of each iCAP occurrence as the number of time points that an iCAP was active or de-active. The average duration, measured in seconds, is the length of time that an iCAP is continuously active. We also evaluated the pair-wise iCAP co-occurrences which is represented by the number of time-points during which a pair of iCAPs were both active divided by the total number of time-points that at least either one of them was active. We also consider the signs of the co-occurrence. Specifically, we considered the pair of iCAPs to be coupled (coupling) if both iCAPs had positive or negative activation or anti-coupled (anti-coupling) if one iCAP was positively/negatively activated and the other had opposite activation (i.e., negatively/positively, respectively). Lastly to measure the temporal overlap between groups of iCAPs, we counted different combinations of iCAPs occurring at each time instance. We then applied hierarchical clustering of iCAPs using the observed combinations as features. Matlab (Mathworks, USA) code for the application of the whole framework can be found at https://www.c4science.ch/source/iCAPs.

### Statistical Analysis

Testing for statistical differences of any iCAP measure were done through a bootstrapping method [73]. A nonparametric approach that makes no distributional assumptions on the observed data. Bootstrapping instead uses resampling to construct empirical confidence intervals (Cis) for a quantity of interest. For each comparison we constructed bootstrap samples by drawing with replacement from the observed measurement. We calculated the null distribution of the observed difference by creating 10,000 bootstrap samples. A 95% CI for the observed difference was obtained by identifying the 2.5^th^ and 97.5^th^ quantiles of the resulting null distribution. The null hypothesis was rejected if 0 was not included in the 95% CI. If more than one comparison was being performed then Bonferroni correction was used. For the analysis of temporal characteristics, we only included the first eight iCAPs as they were equally prevalent across all conditions.

## Data availability

The main data supporting the study are contained within the manuscript. Raw data were too large to be released publicly but are available for research purposes from the corresponding author on reasonable request.

## Code availability

The code for this study is available from the corresponding author on reasonable request.

## Acknowledgments

We thank the NeuroSpin support teams for help in data acquisition and analysis, and Dr. Anjali Tarun and Dr. Daniela Zöller for providing us with human iCAPs for figures and Dr. Younes Farouj for the code to run iCAPs. Part of this work was supported by Institut National de la Santé et de la Recherche Médicale, the Avenir program (B.J.), Commissariat à l’Energie Atomique, Collège de France, ERC Grant NeuroConsc (to S.D.), Foundation Bettencourt-Schueller, the Roger de Spoelberch Foundation, and internal funding from the Department of Physical Medicine and Rehabilitation at the University of Pittsburgh to E.P.

## Author contributions

B.J., S.D., D.V.D.V, E.P. and L.U. designed research; L.U. and B.J. acquired the data; G.H. and J.T. pre-processed the data; S.E. analyzed data; S.E., A.O., and Y.F. adapted the TA framework; B.J., S.D., and E.P. secured funding; and S.E., B.J., and E.P. wrote the paper and all authors contributed to its editing.

