## Supplementary Data for "Transient brain activity dynamics discriminate levels of consciousness during anesthesia"

**Supplementary Figures and Tables**

| **iCAP** | **Lobe** | **Percent** | **Region** | **Z-Score** | **Voxels** |
| --- | --- | --- | --- | --- | --- |
| **1** | Occipital_Lobe (Occipital) | 98.8 | middle_temporal_area (MT) | 2 | 84 |
|  |  | 82.5 | preoccipital_visual_areas_2-3 (V2-V3) | 1.99 | 1807 |
|  |  | 87.1 | visual_area_4 (V4) | 1.99 | 671 |
|  |  | 52.1 | fundus_of_the_superior_temporal_sulcus (STSf) | 1.96 | 232 |
|  |  | 49.6 | primary_visual_cortex (V1) | 1.96 | 952 |
|  |  | 51.7 | ventromedial_intraparietal_sulcus (vm_IPS) | 1.94 | 196 |
|  |  | 96.2 | visual_areas_V6_and_V6A (V6/V6A) | 1.92 | 128 |
|  |  | 95.9 | area_TEO (TEO) | 1.87 | 307 |
|  |  | 20.7 | hippocampal_formation (HF) | 1.85 | 112 |
|  |  | 28.9 | area_7_(PGm)_on_the_medial_wall (area_7m) | 1.81 | 65 |
|  |  | 41.8 | pulvinar_thalamus (Pul) | 1.77 | 81 |
|  |  | 86.3 | medial_superior_temporal_area (MST) | 1.76 | 126 |
| **iCAP** | **Lobe** | **Percent** | **Region** | **Z-Score** | **Voxels** |
| **2** | telencephalon (tel) | 100 | secondary_somatosensory_cortex (SII) | 2.67 | 322 |
|  |  | 97 | floor_of_the_lateral_sulcus (floor_of_ls) | 2.6 | 262 |
|  |  | 83.3 | claustrum (Cl) | 2.36 | 250 |
|  |  | 53 | dorsal_striatum (DStr) | 2.24 | 1126 |
|  |  | 80.6 | caudal_orbital_frontal_cortex (caudal_OFC) | 2.2 | 325 |
|  |  | 26.3 | lateral_motor_cortex (M1/PM) | 2.09 | 593 |
|  |  | 30.3 | primary_somatosensory_cortex (SI) | 2.01 | 282 |
|  |  | 82.4 | core_areas_of_auditory_cortex (core) | 1.96 | 145 |
|  |  | 24.2 | area_7_in_the_inferior_parietal_lobule (area_7_in_IPL) | 1.89 | 157 |
|  |  | 68.9 | belt_areas_of_auditory_cortex (belt) | 1.88 | 219 |
|  |  | 27.2 | caudal_superior_temporal_gyrus (STGc) | 1.62 | 88 |
|  |  | 36.2 | septum_diagonal_band_complex (SDBR) | 1.61 | 42 |
|  |  | 23.9 | rostral_superior_temporal_region (STGr/STSd) | 1.61 | 164 |
| **iCAP** | **Lobe** | **Percent** | **Region** | **Z-Score** | **Voxels** |
| **3** | Frontal_Lobe (Frontal); Parietal_Lobe (Parietal) | 72.5 | primary_somatosensory_cortex (SI) | 2.63 | 676 |
|  |  | 100 | midcingulate_cortex (MCC) | 2.47 | 207 |
|  |  | 73.5 | lateral_motor_cortex (M1/PM) | 2.42 | 1655 |
|  |  | 81.3 | area_5 (area_5) | 2.29 | 335 |
|  |  | 90.9 | medial_supplementary_motor_areas (SMA/preSMA) | 2.11 | 319 |
|  |  | 52.6 | lateral_intraparietal_sulcus (lat_IPS) | 2 | 205 |
|  |  | 32.4 | area_7_(PGm)_on_the_medial_wall (area_7m) | 1.98 | 73 |
|  |  | 52.3 | posterior_cingulate_gyrus (PCgG) | 1.97 | 320 |
|  |  | 24.2 | anterior_cingulate_cortex (ACC) | 1.93 | 126 |
|  |  | 38.9 | area_7_in_the_inferior_parietal_lobule (area_7_in_IPL) | 1.82 | 252 |
|  |  | 65.7 | periarcuate_area_8A_(Frontal_Eye_Fields) (area_8A) | 1.76 | 44 |
| **iCAP** | **Lobe** | **Percent** | **Region** | **Z-Score** | **Voxels** |
| **4** | Frontal_Lobe (Frontal); telencephalon (tel) | 100 | medial_orbital_frontal_cortex (med_OFC) | 2.37 | 294 |
|  |  | 98.7 | anterior_cingulate_cortex (ACC) | 2.35 | 514 |
|  |  | 99.4 | lateral_orbital_frontal_cortex (lat_OFC) | 2.31 | 702 |
|  |  | 94 | ventrolateral_prefrontal_cortex (vlPFC) | 2.02 | 671 |
|  |  | 53.5 | dorsal_striatum (DStr) | 1.93 | 1137 |
|  |  | 61 | medial_supplementary_motor_areas (SMA/preSMA) | 1.89 | 214 |
|  |  | 79.5 | dorsolateral_prefrontal_cortex (dlPFC) | 1.88 | 843 |
|  |  | 35.2 | ventral_striatum (VStr) | 1.8 | 45 |
|  |  | 59.3 | caudal_orbital_frontal_cortex (caudal_OFC) | 1.72 | 239 |
|  |  | 23.7 | midcingulate_cortex (MCC) | 1.67 | 49 |
| **iCAP** | **Lobe** | **Percent** | **Region** | **Z-Score** | **Voxels** |
| **5** | Temporal_Lobe (Temporal); telencephalon (tel) | 99.2 | rhinal_cortex (Rh) | 2.91 | 386 |
|  |  | 100 | lateropallial_amygdala (lpAmy) | 2.82 | 118 |
|  |  | 96.4 | temporal_pole (TG) | 2.68 | 406 |
|  |  | 100 | ventropallial_amygdala (vpAmy) | 2.57 | 42 |
|  |  | 65 | piriform_cortex (Pir) | 2.34 | 26 |
|  |  | 95.8 | area_TE (TE) | 2.33 | 1059 |
|  |  | 100 | endopiriform_claustrum (En) | 2.3 | 32 |
|  |  | 78.2 | hippocampal_formation (HF) | 2.27 | 423 |
|  |  | 61.9 | rostral_superior_temporal_region (STGr/STSd) | 2.08 | 425 |
|  |  | 41.6 | fundus_of_the_superior_temporal_sulcus (STSf) | 2.07 | 185 |
|  |  | 30 | claustrum (Cl) | 2.05 | 90 |
|  |  | 91.5 | parahippocampal_cortex (paraHipp) | 2.05 | 129 |
|  |  | 63.6 | medial_amygdala (mAmy) | 1.89 | 35 |
|  |  | 29.5 | floculus_parafloculus (Fl-PFl) | 1.62 | 52 |
| **iCAP** | **Lobe** | **Percent** | **Region** | **Z-Score** | **Voxels** |
| **6** | myelencephalon (myel); metencephalon (met); mesencephalon (mes) | 100 | deep_cerebellar_nuclei (DCb) | 3.1 | 40 |
|  |  | 100 | parabrachial_complex (PBC) | 2.44 | 80 |
|  |  | 100 | intermediate_cerebellar_cortex (ICbCx) | 2.43 | 1178 |
|  |  | 96.8 | vermis_cerebellar_cortex (VCbCx) | 2.24 | 1331 |
|  |  | 99.6 | pontine_nucleus_region (Pn+) | 2.21 | 268 |
|  |  | 93.8 | floculus_parafloculus (Fl-PFl) | 2.16 | 165 |
|  |  | 100 | midbrain_tegmentum (TgMid) | 2.14 | 47 |
|  |  | 100 | pontine_reticulum (retPons) | 2.09 | 71 |
|  |  | 91.1 | lateral_cerebellar_cortex (LCbCx) | 2.04 | 350 |
|  |  | 57.5 | colliculi (Co) | 1.91 | 69 |
|  |  | 89.3 | periaqueductal_gray_region (PAGR) | 1.85 | 75 |
|  |  | 61.3 | midbrain_reticulum (RtMid) | 1.78 | 49 |
|  |  | 22.4 | preoccipital_visual_areas_2-3 (V2-V3) | 1.76 | 491 |
|  |  | 55.2 | midbrain_dopaminergic_complex (DA_Mid) | 1.74 | 133 |
| **iCAP** | **Lobe** | **Percent** | **Region** | **Z-Score** | **Voxels** |
| **7** | Parietal_Lobe (Parietal) | 100 | area_7_(PGm)_on_the_medial_wall (area_7m) | 3.17 | 225 |
|  |  | 94.6 | lateral_intraparietal_sulcus (lat_IPS) | 2.86 | 369 |
|  |  | 89.6 | area_5 (area_5) | 2.86 | 369 |
|  |  | 74.8 | area_7_in_the_inferior_parietal_lobule (area_7_in_IPL) | 2.83 | 485 |
|  |  | 97.1 | ventromedial_intraparietal_sulcus (vm_IPS) | 2.82 | 368 |
|  |  | 33.9 | visual_area_4 (V4) | 2.6 | 261 |
|  |  | 68.4 | visual_areas_V6_and_V6A (V6/V6A) | 2.45 | 91 |
|  |  | 92.5 | medial_superior_temporal_area (MST) | 2.29 | 135 |
|  |  | 65.9 | middle_temporal_area (MT) | 2.19 | 56 |
|  |  | 24.8 | caudal_superior_temporal_gyrus (STGc) | 2.17 | 80 |
|  |  | 72.4 | posterior_cingulate_gyrus (PCgG) | 2.16 | 443 |
|  |  | 20.9 | preoccipital_visual_areas_2-3 (V2-V3) | 2.02 | 458 |
| **iCAP** | **Lobe** | **Percent** | **Region** | **Z-Score** | **Voxels** |
| **8** | Frontal_Lobe (Frontal) | 97.4 | dorsolateral_prefrontal_cortex (dlPFC) | 3.84 | 1033 |
|  |  | 36.1 | medial_orbital_frontal_cortex (med_OFC) | 2.59 | 106 |
|  |  | 29 | lateral_motor_cortex (M1/PM) | 2.55 | 653 |
|  |  | 71.8 | medial_supplementary_motor_areas (SMA/preSMA) | 2.42 | 252 |
|  |  | 66.8 | ventrolateral_prefrontal_cortex (vlPFC) | 2.41 | 477 |
|  |  | 81.2 | anterior_cingulate_cortex (ACC) | 2.16 | 423 |
| **iCAP** | **Lobe** | **Percent** | **Region** | **Z-Score** | **Voxels** |
| **9** | Occipital_Lobe (Occipital) | 61.1 | primary_visual_cortex (V1) | 3.51 | 1171 |
|  |  | 25.2 | preoccipital_visual_areas_2-3 (V2-V3) | 2.48 | 553 |
|  |  | 24.9 | vermis_cerebellar_cortex (VCbCx) | 2.04 | 343 |
| **iCAP** | **Lobe** | **Percent** | **Region** | **Z-Score** | **Voxels** |
| **10** | Occipital_Lobe (Occipital) | 47 | primary_visual_cortex (V1) | 3.9 | 901 |
|  |  | 20.3 | preoccipital_visual_areas_2-3 (V2-V3) | 2.8 | 444 |
|  |  | 70.7 | visual_areas_V6_and_V6A (V6/V6A) | 2.05 | 94 |
| **iCAP** | **Lobe** | **Percent** | **Region** | **Z-Score** | **Voxels** |
| **11** | metencephalon (met) | 42.9 | vermis_cerebellar_cortex (VCbCx) | 4.56 | 590 |
|  |  | 27.4 | intermediate_cerebellar_cortex (ICbCx) | 4.06 | 323 |
|  |  | 37.8 | lateral_cerebellar_cortex (LCbCx) | 4.06 | 145 |

**Supplementary Table 1**: We compute the average percent coverage, z-score, and the total number of voxels occupied in brain areas defined with the CHARM and SARM atlas [26-28].

|  | Awake | Ketamine | Moderate Sevoflurane | Deep Sevoflurane | Moderate Propofol | Deep Propofol | Totals |
| --- | --- | --- | --- | --- | --- | --- | --- |
| MK-J | 18 | 0 | 5 | 2 | 1 | 6 | 32 |
| MK-Ki | 0 | 7 | 10 | 8 | 0 | 0 | 25 |
| MK-A | 4 | 0 | 0 | 0 | 0 | 0 | 4 |
| MK-K | 9 | 5 | 0 | 0 | 7 | 9 | 30 |
| MK-R | 0 | 10 | 8 | 10 | 12 | 12 | 52 |
| Totals | 31 | 22 | 23 | 20 | 20 | 27 | 143 |

**Supplementary Table 2**: Number of sessions for each monkey and for awake and each type of anesthetic.

| **Paper** | Karahanoglu 2015 | Zoeller 2018 | Zoeller 2019 | Zoeller 2021 | Piguet 2021 | Tarun 2020 | Pirondini 2022 |
| --- | --- | --- | --- | --- | --- | --- | --- |
| **TR** | 1.1 | 2.4 | 2.4 | 2.4 | 2.1 | 2.1 | 2 |
| **Number of iCAPs** | 20 | 18 | 17 | 17 | 20 | 17 | 16 |
| **Auditory** |  |  |  |  |  |  |  |
| **Language** |  |  |  |  |  |  |  |
| **Executive** | anterior |  |  |  |  | left  /right |  |
| **Attention/**  **FPN** |  | left/  right |  |  |  |  |  |
| **Primary Visual** |  |  | 2 | 2 | 3 |  | 3 |
| **Secondary Visual/Higher Visual** |  |  |  |  |  |  |  |
| **DMN** |  |  |  |  |  |  |  |
| **aDMN/ACC** |  |  |  |  |  |  |  |
| **pDMN** |  |  |  |  |  |  |  |
| **Salience** | anterior |  | anterior | anterior | anterior/full (2) |  | anterior |
| **OFC** |  |  |  |  |  |  |  |
| **Precuneus/**  **PCC/Thalamus** |  |  |  |  |  |  |  |
| **CEB** |  |  |  |  |  |  |  |
| **aCEB** |  |  |  |  |  |  |  |
| **pCEB** |  |  |  |  |  |  |  |
| **Visuospatial/**  **motor/**  **sensory** |  | 3 |  |  |  | 3 |  |
| **Sensorimotor** |  | 2 |  |  |  |  |  |
| **Temporal/**  **Amygdala/**  **iTEMP/FUS** |  |  | 2 | 2 |  |  |  |
| **PCC/Thalamus** |  |  |  |  |  |  |  |
| **Subcortical** |  |  |  |  |  |  |  |
| **Insula** |  | anterior |  |  |  |  |  |
| **Prefrontal** |  |  |  |  |  |  |  |

**Supplementary Table 3**: iCAP Studies used to compare human spatial iCAPs to monkey spatial iCAPs. For each study, green color indicates if iCAP was present and number indicates how many of that cluster was present.

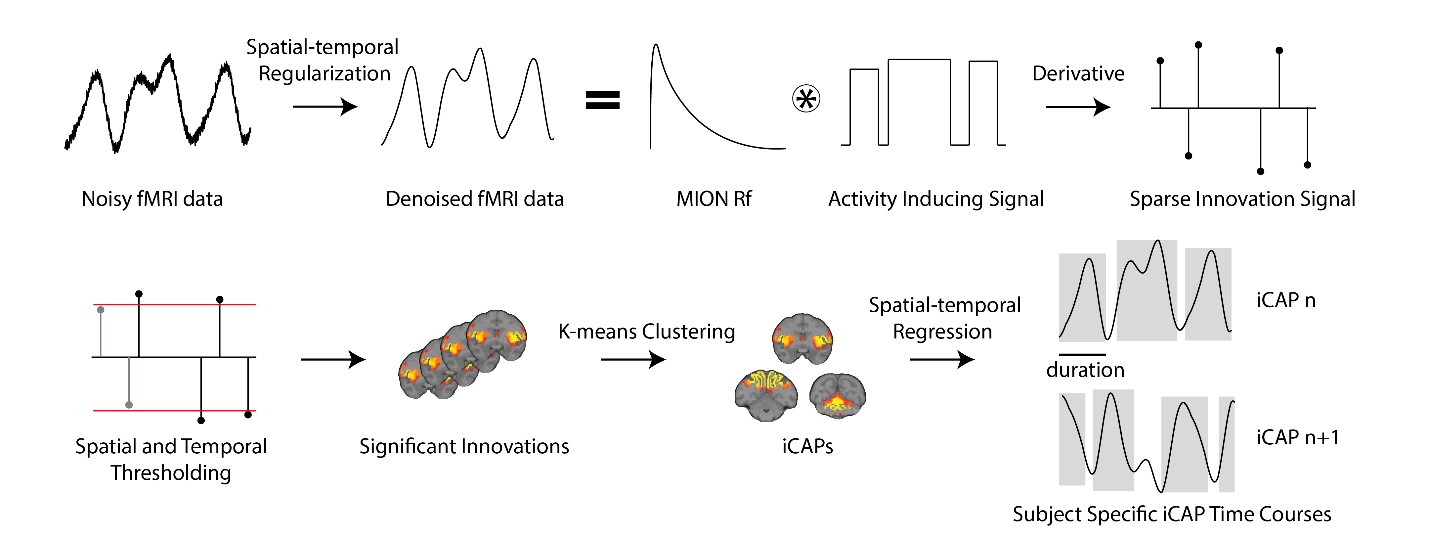

**Supplementary Fig. 1 | Methodological pipeline of the iCAP framework.** **A)** Noisy fMRI timecourses are denoised using a combined spatio-temporal regression, followed by a deconvolution from the MION rf to obtain a block type activity inducing signal, which is then differentiated to get the sparse innovation signal. The resulting innovation signals undergo a two-step spatial and temporal thresholding to select significant innovation frames. Then, the latter undergo a k-means clustering to obtain stable large-scale networks (i.e. iCAPs). The iCAPs are then back projected to the individual activity inducing signals using spatial-temporal regression to recover temporal profiles of each iCAP.

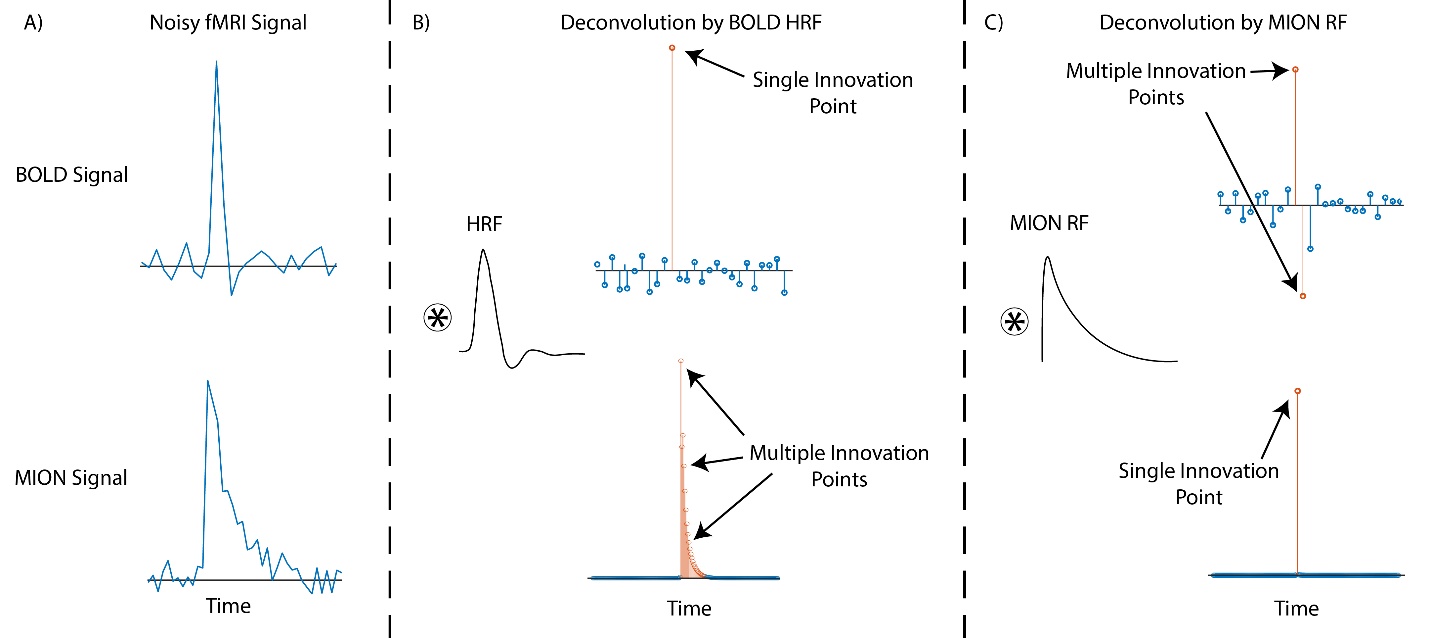

**Supplementary Fig. 2 | BOLD HRF vs MION RF. A)** Left column is representative of a noisy fMRI signal measured using BOLD and MION, respectively top and bottom. **B)** Middle column is the noisy fMRI signals deconvolved using the canonical BOLD HRF. **C)** Right column is the noisy fMRI signals deconvolved using the MION RF. It can be seen that the MION RF gets a much cleaner deconvolution of the MION signal (i.e., a single time innovation point) as compared to the BOLD HRF. Instead, the BOLD HRF gets a much cleaner deconvolution of the BOLD signal (i.e. a single time innovation point) as compared to the MION RF.

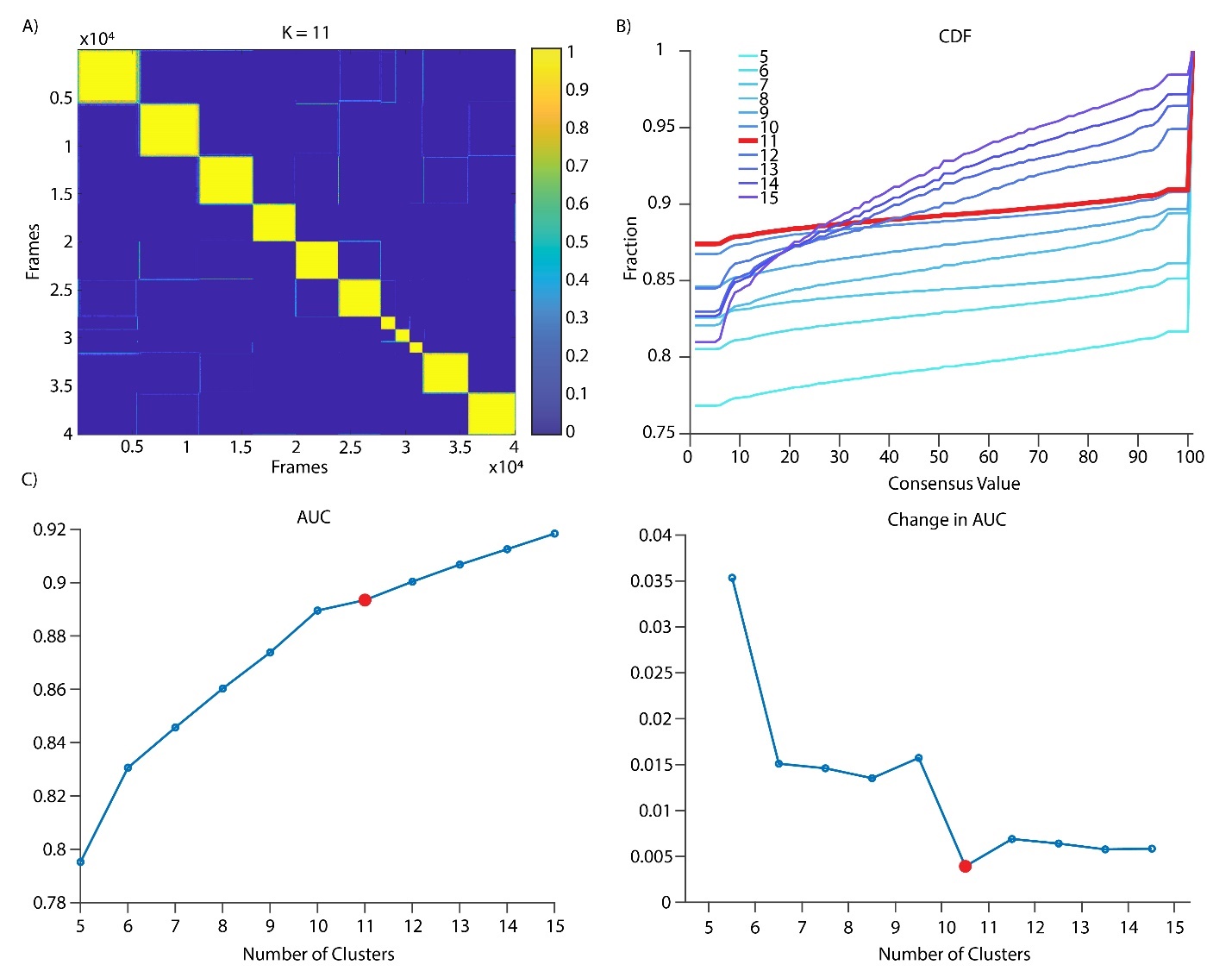

**Supplementary Fig. 3 | Optimal Clustering. A)** Consensus clustering matrix for n = 11 clusters. The x and y axes correspond to frame number. Values in the matrix range from 0 to 1, which indicate the reproducibility of the sampling across multiple runs, with 1 being perfectly re-sampled at all times. Diagonal values are expected to be equal to 1 (the same frame indices will always be clustered into the same group). **B)** The cumulative distribution function (CDF) indicates the extent to which the consensus matrix distribution is skewed toward 0 and 1, with a flat line being the ideal shape (i.e. 0 means two frames are never clustered together while 1 means frames are always clustered together). In red CDF for n = 11 clusters. **C)** The area under the curve (AUC) of the CDF and the change in AUC display the optimal number of cluster K to which there is minimal increase in the AUC. In red the values for n = 11 clusters.

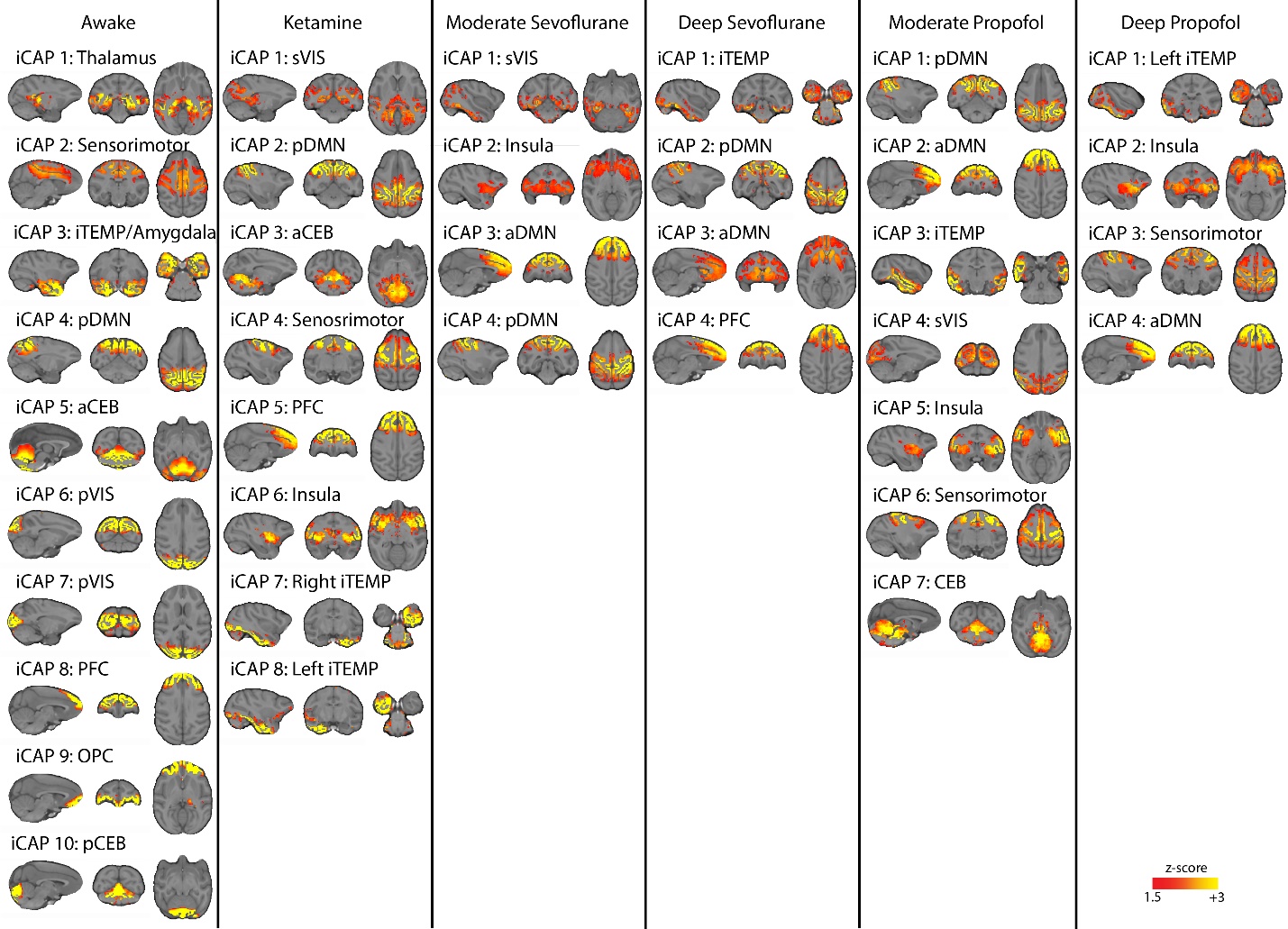

**Supplementary Fig. 4 | Spatial iCAPs for Each Condition.** Spatial pattern of the optimal iCAPs found when clustering significant innovation frames for each condition separately.
